## Supplementary material for "Extensive topographic remapping and functional sharpening in the adult rat visual pathway upon first visual experience": figs and tables single document

**This file includes:**

S1 Text
Figs A-I
Tables A-E

### S1 Text

#### 1. Setup

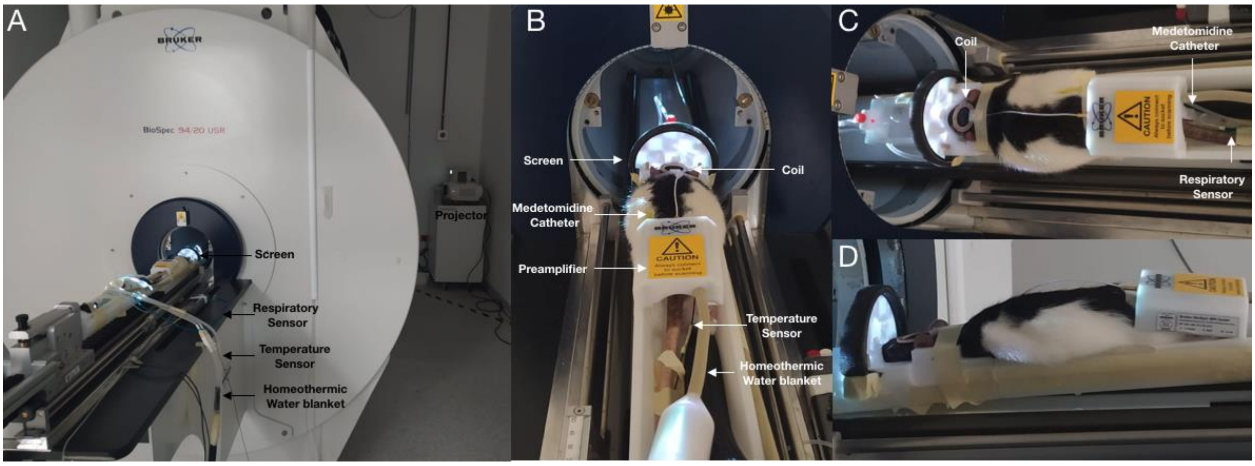


**Fig A. Photographs of the visual setup mounted to an MRI animal cradle.** Animal life-support equipment: temperature and respiration sensors, subcutaneous catheter, and homeothermic water blanket were connected to the cradle. The surface coil was placed on the head of the rats and the preamplifier was placed behind the animal. The visual stimulus was projected to a mirror that reflected the image to the screen placed in front of the animal eyes.

#### 2. Robust activation upon complex stimulation

Fig B shows that the retinotopic and SF tuning stimuli elicited reliable and robust BOLD activation throughout the entire visual pathway of HC, i.e. LGN, VC and SC. Panels A and E in Fig B show the percentage of BOLD signal change (PSC) of the LGN, VC and SC (Panel D in Fig B) averaged across animals, runs and cycles obtained for both types of complex stimuli. The SF tuning response is overall stronger than the response to the retinotopy stimulus, likely driven by the fact that the SF tuning stimulus is observed across the entire field of view and that the contrast of SF tuning stimulus (100%) was double that of the retinotopy stimulus.

The retinotopy stimulus yielded stronger BOLD responses to the vertical movement, i.e. in the second (84-118 TRs) and fourth (192-216 TRs) stimulation blocks, compared to the horizontal movement (in the first (30-54 TRs) and third (138-162 TRs) blocks), likely due to the preference for vertical movements compared to horizontal ones in rats [1].

In both stimuli, SC has the strongest response, followed by LGN, and VC has the lowest BOLD response amplitude, following the hierarchical preprocessing from early order areas (LGN, SC) to later areas in the processing of visual information (VC). Also, all visual structures show a post-stimulus overshoot, most noticeable in SC.

Importantly, the activation maps show robust responses limited to the areas of the visual pathway. The activation patterns are consistent across scanning sessions, in particular with the retinotopy stimulus (Fig 4 and Fig C in S1 Text). For the SF tuning stimulus, the responses of LGN and SC are consistent across scanning sessions, however, the VC's response at t=17d and t=27d is absent or highly attenuated when compared to t=0 and t=7d. This suggests an adaptation to these strong stimuli, rather than e.g. loss of sensitivity given that the VC's response to the retinotopy stimulus remains highly robust at all the time points measured.

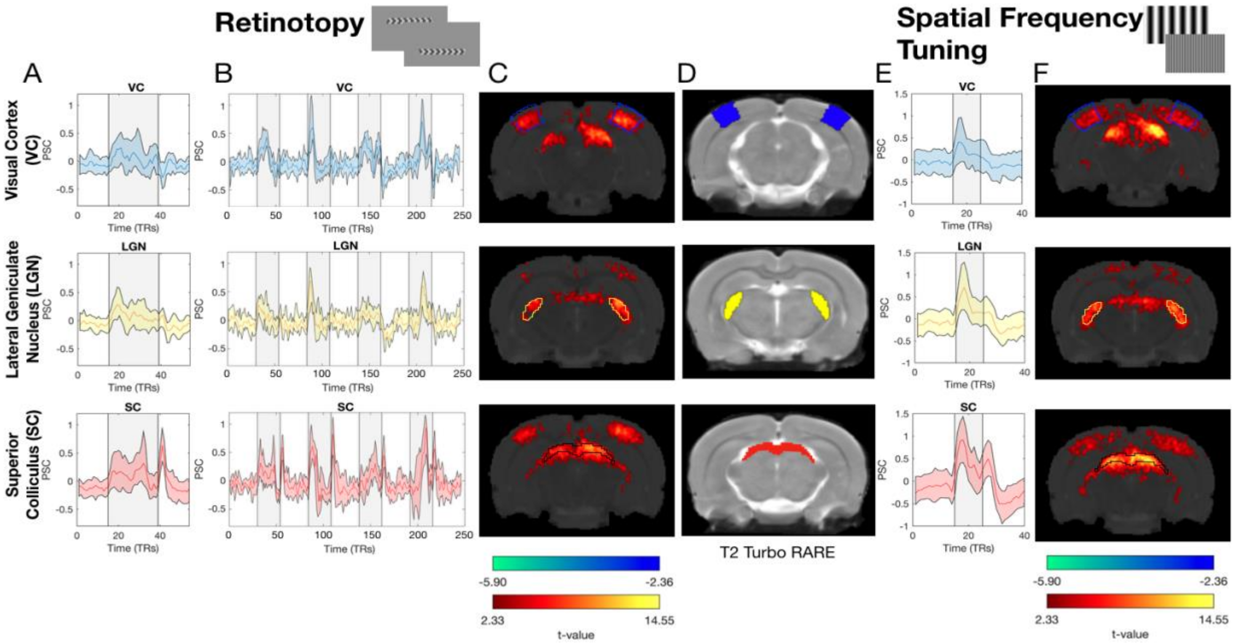


**Fig B. Retinotopic and SF tuning visual stimuli result in robust BOLD signals confined to the visual pathway in HC at t=0.** A, B and E: Percentage of BOLD signal change (PSC) of the ROIs defined in D averaged across animals, runs (B) and cycles (A, E), upon retinotopic (A, B) and SF tuning (E) visual stimulation. The colored areas correspond to the 95% confidence interval and the gray area to the stimulation period. C and F: GLM functional maps obtained after retinotopic (C) and SF tuning (F) visual stimulation. The maps are FDR corrected using a p-value of 0.001 and minimum cluster size of 20 voxels. The ROIs defined based on the SIGMA atlas are overlaid on the functional maps. D: Anatomical images with the delineation of the ROIs. The data underlying this figure can be found here: doi:10.18112/openneuro.ds004509.v1.0.0.

3. Differential BOLD responses to the SF tuning stimulus between VD and HC

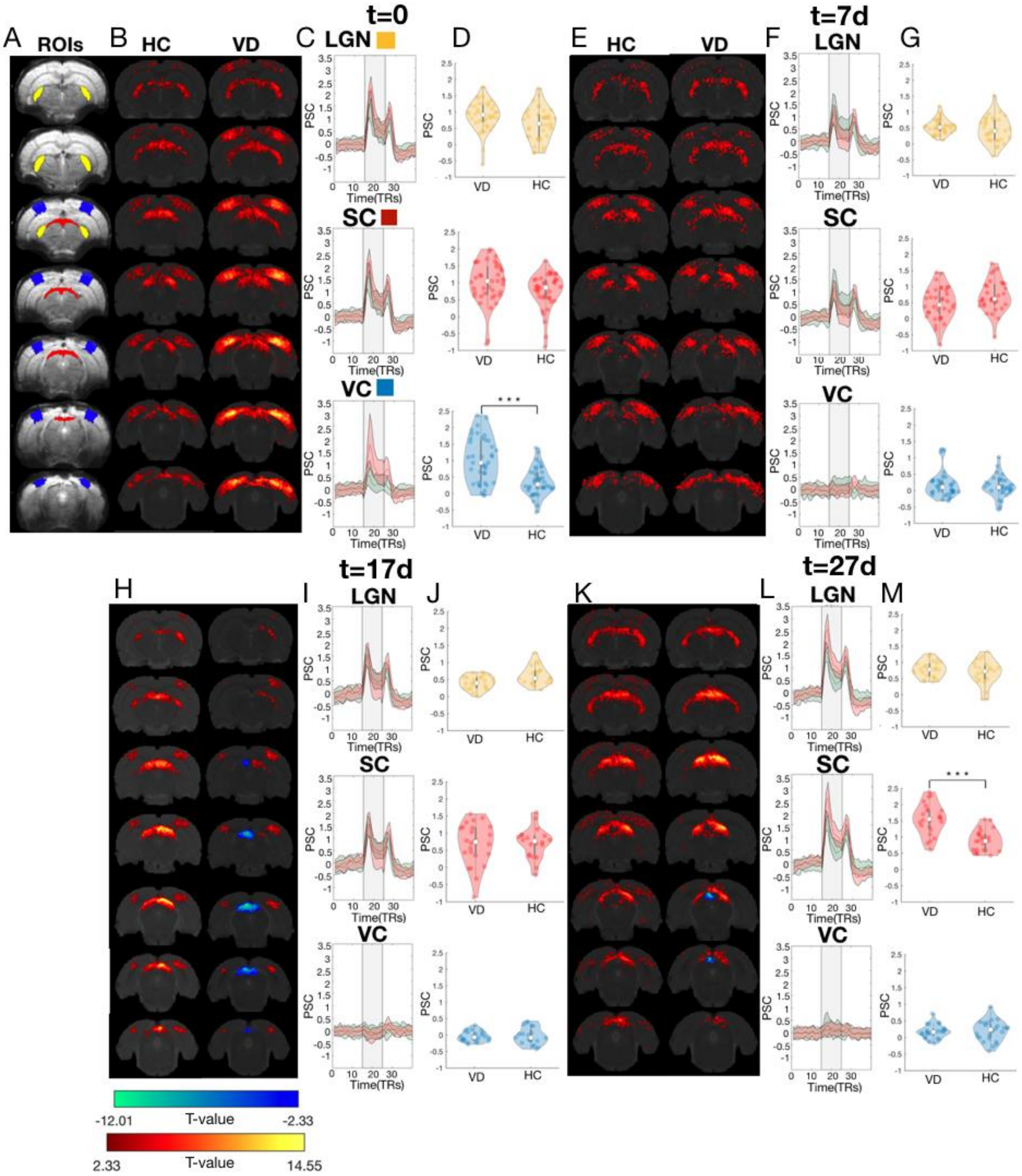


**Fig C. Differential responses between VD animals and HC driven by the SF tuning stimulus.** A: Raw fMRI images with the ROIs (LGN, SC and VC) overlaid. B, E, H, K: fMRI activation patterns of t-contrast maps obtained for HC and VD animals at t=0, t=7d, t=17d and t=27d, respectively. The GLM maps are FDR corrected using a p-value of 0.001 and minimum cluster size of 20 voxels. C, F, I, L: PSC of the LGN, SC and VC for the HC (green) and VD (red) animals at t=0, t=7d, t=17d and t=27d, respectively. The grey area represents the stimulation period. D, G, J, M: Violin plot of the amplitude of the BOLD response of VD and HC during the total duration of the activation period obtained with the SF tuning stimulus at t=0, t=7d, t=17d and t=27d, respectively. The white dot represents the mean, and the grey bar represents the 25% and 75% percentiles. The yellow, red and blue colors represent the LGN, SC and VC respectively. The *** represents a p-value<0.001, ** p-value<0.01 and * p-value<0.05. The data underlying this figure can be found here: doi:10.18112/openneuro.ds004509.v1.0.0.

### 4. Differential BOLD responses to the retinotopic stimulus between VD and HC

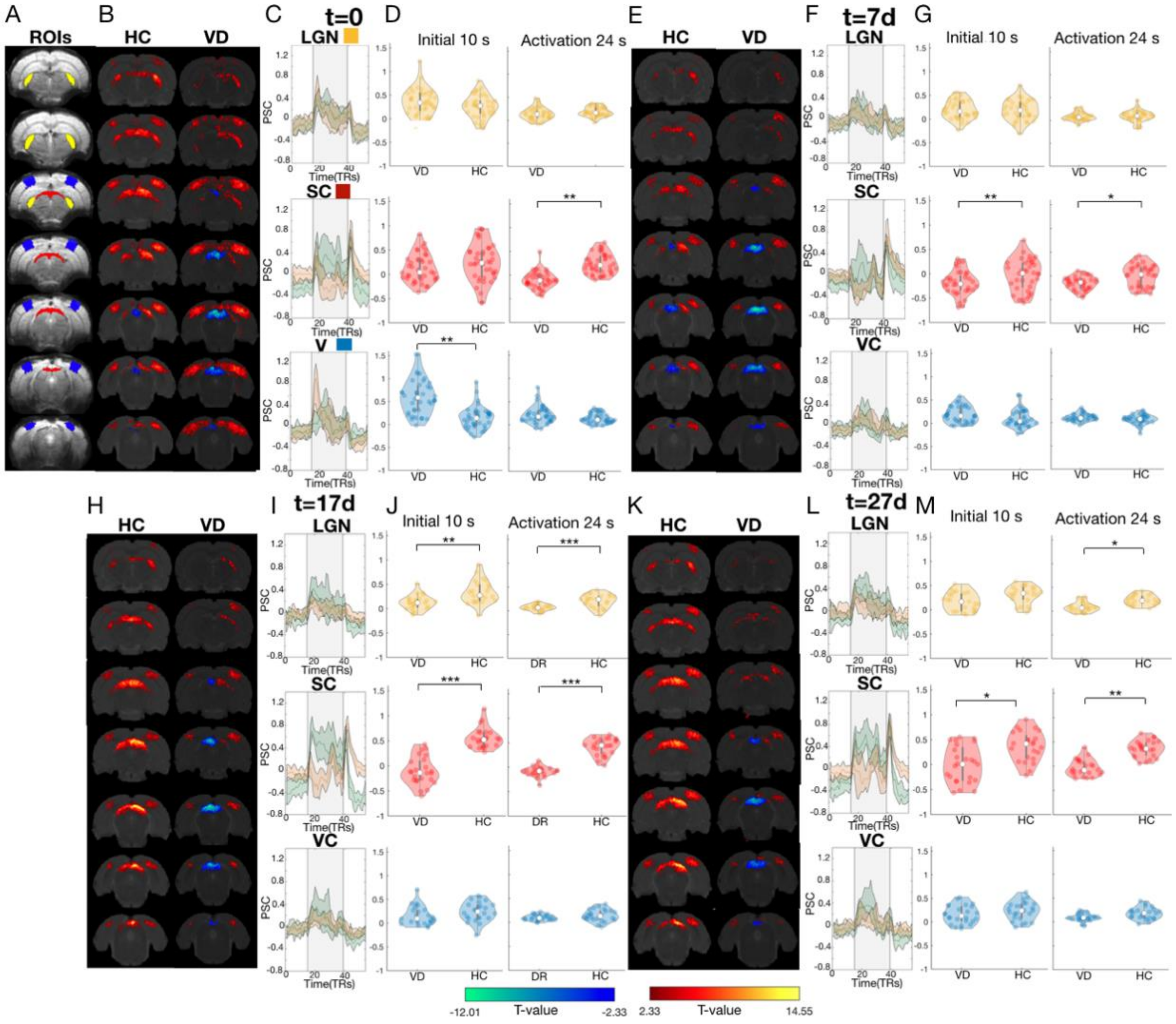


**Fig D. Differential responses between VD animals and HC driven by the retinotopic stimulus.** Similar Figure to Fig4, but the violin plots now show the amplitude of the BOLD response during each stimulation block. A: Raw fMRI images with the ROIs (LGN, SC and VC) overlaid. B, E, H, K: fMRI activation patterns of t-contrast maps obtained for HC and VD animals at t=0, t=7d, t=17d and t=27d, respectively. The GLM maps are FDR corrected using a p-value of 0.001 and minimum cluster size of 20 voxels. C, F, I, L: PSC of the LGN, SC and VC for the HC (green) and VD (orange) animals at t=0, t=7d, t=17d and t=27d, respectively. The grey area represents the stimulation period. D, G, J, M: Violin plot of the amplitude of the BOLD response of VD and HC during the initial 10 s of the activation period (left) and the total duration of the activation period obtained with the retinotopy stimulus (right) at t=0, t=7d, t=17d and t=27d, respectively. Each dot represents the average BOLD response during a stimulation block. The white dot represents the mean, and the grey bar represents the 25% and 75% percentiles. The yellow, red and blue colors represent the LGN, SC and VC respectively. The *** represents a p-value<0.001, ** p-value<0.01 and * p-value<0.05. The data underlying this figure can be found here: doi:10.18112/openneuro.ds004509.v1.0.0.

#### 4.1. BOLD statistical analysis

| Initial 10 s | Activation 24 s |
|---|---|

| | VC | LGN | SC | VC | LGN | SC |
|---|---|---|---|---|---|---|
| t=0 | **0.003** | 0.400 | 0.110 | 0.750 | 0.16 | **0.003** |
| t=7d | 0.120 | 0.850 | **0.01** | 0.240 | 0.760 | **0.03** |
| t=17d | 0.100 | **0.008** | **$7x10^{-4}$** | 0.090 | **$8x10^{-4}$** | **$2.2x10^{-5}$** |
| t=27d | 0.300 | 0.055 | **0.018** | 0.130 | **0.025** | **0.01** |

**Table A. Statistical analysis of HC vs VD BOLD changes.** P-values associated with the ANOVA Bonferroni corrected for multiple comparisons (brain areas and sessions) statistical analysis of the BOLD amplitude changes between HC and VD in response to the retinotopic stimulus (Fig 4) during the initial 10 s of the activation period and the total duration of the activation period. P-values below 0.05 are shown in bold. The data underlying this table can be found here: doi:10.18112/openneuro.ds004509.v1.0.0.

#### 4.2. PRF size statistical analysis

| Brain area | Session | Group | Contrast | p-value |
|---|---|---|---|---|
| **VC** | **t=0** | | **HC vs VD** | **0.045** |
| LGN | t=0 | | HC vs VD | 0.085 |
| **SC** | **t=0** | | **HC vs VD** | **0.039** |
| VC | t=7d | | HC vs VD | 1.000 |
| LGN | t=7d | | HC vs VD | 0.889 |
| SC | t=7d | | HC vs VD | 0.998 |
| VC | t=17d | | HC vs VD | 1.000 |
| **LGN** | **t=17d** | | **HC vs VD** | **0.023** |
| SC | t=17d | | HC vs VD | 0.582 |
| VC | t=27d | | HC vs VD | 0.948 |
| **LGN** | **t=27d** | | **HC vs VD** | **0.018** |
| SC | t=27d | | HC vs VD | 0.998 |
| VC | | HC | t=0 vs t=7d | 0.985 |
| VC | | HC | t=7d vs t=17d | 1.000 |
| VC | | HC | t=17d vs t=27d | 1.000 |
| **LGN** | | **HC** | **t=0 vs t=7d** | **0.04** |
| LGN | | HC | t=7d vs t=17d | 0.936 |
| LGN | | HC | t=17d vs t=27d | 0.943 |

| | | | | |
|---|---|---|---|---|
| SC | | HC | t=0 vs t=7d | 0.777 |
| SC | | HC | t=7d vs t=17d | 0.775 |
| SC | | HC | t=17d vs t=27d | 1.000 |
| **VC** | | **VD** | **t=0 vs t=7d** | **0.048** |
| VC | | VD | t=7d vs t=17d | 1.000 |
| VC | | VD | t=17d vs t=27d | 0.999 |
| **LGN** | | **VD** | **t=0 vs t=7d** | **0.041** |
| LGN | | VD | t=7d vs t=17d | 1.000 |
| LGN | | VD | t=17d vs t=27d | 1.000 |
| **SC** | | **VD** | **t=0 vs t=7d** | **0.043** |
| SC | | VD | t=7d vs t=17d | 1.000 |
| SC | | VD | t=17d vs t=27d | 1.000 |

**Table B. P-values obtained for the pRF size changes between HC and VD, calculated using ANOVA Bonferroni corrected for multiple comparisons (brain areas and sessions).** P-values below 0.05 are shown in bold. The data underlying this table can be found here: doi:10.18112/openneuro.ds004509.v1.0.0.

#### 5. Literature review on the optimal spatial frequency

| Study | Visual area | Optimal SF | Species | Technique |
|---|---|---|---|---|
| [2] | VC | 0.036 cpd | Mouse | Electrophysiology |
| [3] | VC | 0.05 cpd | Mouse | Electrophysiology |
| [4] | VC | 0.03 cpd | Mouse | Electrophysiology |
| [5] | VC | 0.04 cpd | Mouse | Electrophysiology |
| [6] | VC | 0.046 cpd | Mouse | Electrophysiology<br>Intrinsic Signal Optical Imaging |
| [7] | VC | 0.08cpd | Mouse | Two Photon Imaging |
| [8] | VC | 0.031 cpd | Mouse | Electrophysiology |
| [9] | VC | 0.1 cpd | Rat | Electrophysiology |
| [10] | VC | 0.05-0.06 cpd | Rat | Electrophysiology |
| [11] | VC | 0.068 cpd | Rat | Electrophysiology |
| [12] | LGN | 0.027 cpd | Mouse | Electrophysiology |

| | | | | |
|---|---|---|---|---|
| [3] | LGN | 0.08 cpd | Mouse | Electrophysiology |
| [13] | LGN | 0.09 cpd | Mouse | Electrophysiology |
| [14] | LGN | 0.02 - 0.05 cpd | Mouse | Electrophysiology |
| [15] | LGN | 0.035 cpd | Mouse | Electrophysiology |
| [16] | LGN | 0.03-0.06 cpd | Rat | Electrophysiology |
| [17] | LGN | 0.04-0.05 cpd | Rat | Electrophysiology |
| [18] | SC | 0.08 cpd | Mouse | Electrophysiology |
| [19] | SC | 0.048 cpd | Mouse | Electrophysiology |
| [20] | SC | 0.1 cpd | Mouse | Electrophysiology |
| [21] | SC | 0.03 cpd | Rat | Electrophysiology |
| [11] | SC | 0.03 cpd | Rat | Electrophysiology |

**Table C. Summary of the optimal spatial frequency measured for rat and mice across multiple studies.** The rat studies used to define the reference interval of Figure 3 are highlighted in yellow. The data underlying this table can be found here: doi:10.18112/openneuro.ds004509.v1.0.0.

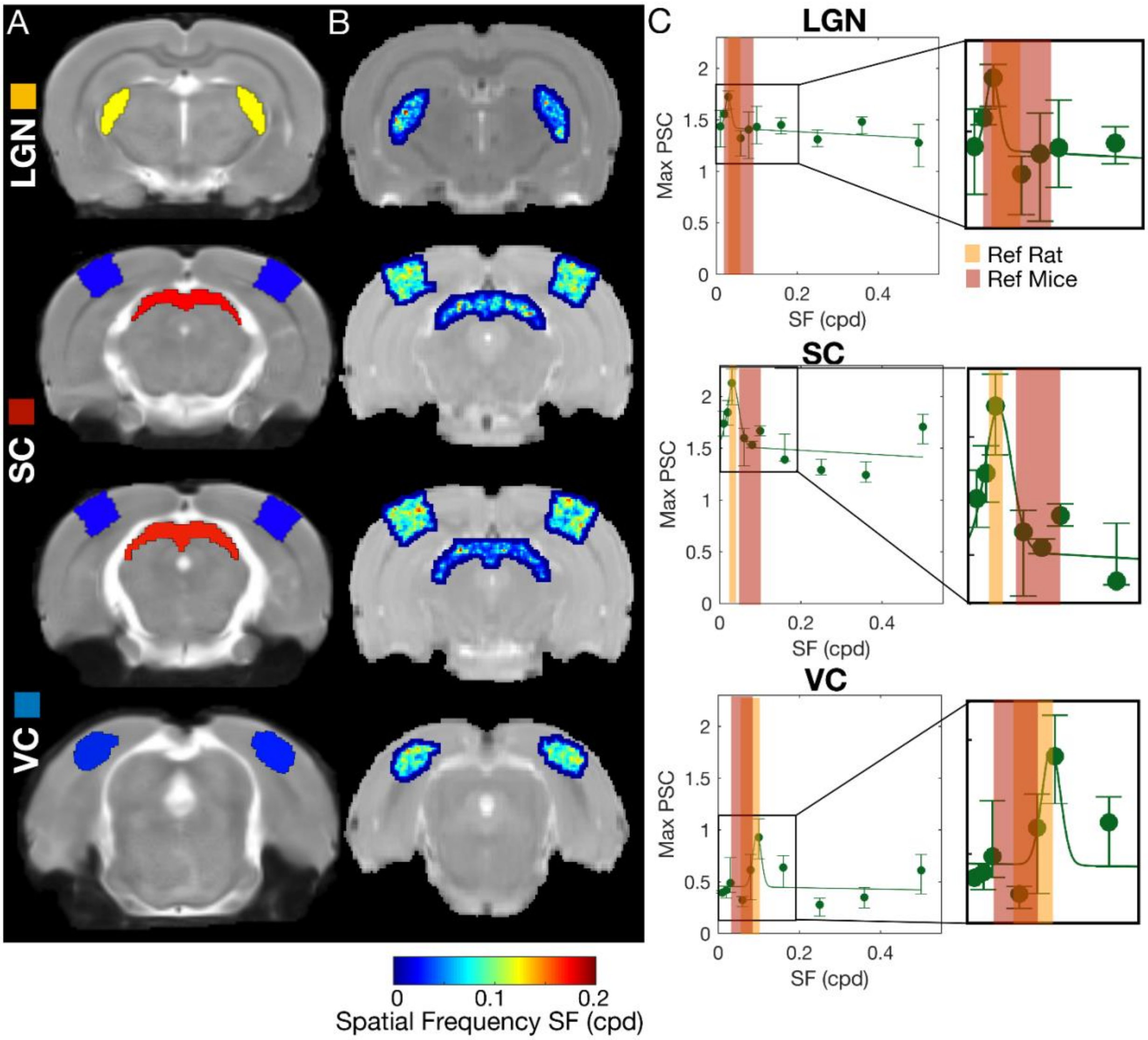


**Fig E. Spatial frequency selectivity across the visual pathway in HC animals at t=0.** A: Anatomical images with the ROIs highlighted. B: Optimal spatial frequency estimated per voxel for HC, averaged across animals. C: Maximum PSC during the activation period as a function of the spatial frequency of the stimulus, calculated for HC. The errorbar represents the 10% confidence interval across animals. The continuous lines represent the Gaussian model fitted to the data. The goodness of fit is shown in Table D. The orange and red bands denote the range of optimal spatial frequency values reported in the literature measured using electrophysiology for rats and mice, respectively. A compilation of 22 studies reporting on the optimal spatial frequency of the rat and mouse visual pathway can be found in Table C. The data underlying this figure can be found here: doi:10.18112/openneuro.ds004509.v1.0.0.

#### 6. Quantification of the retinotopic organization of the visual pathway

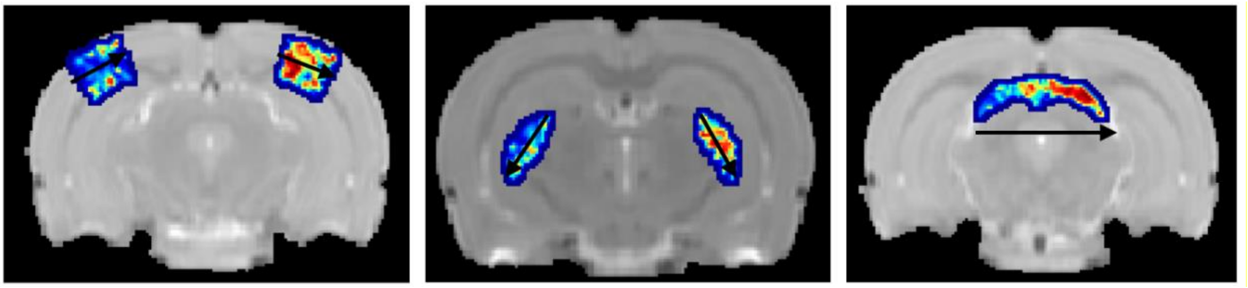

**Fig F. Gradient direction for VC (left), LGN (middle) and SC (right).**

We applied a complementary approach to quantify the retinotopic arrangement of the visual pathway across scanning sessions. This approach does not only take into account the pRF phase but also its eccentricity and size. Here we looked at the pRF profiles estimated with micro probing technique (Panel A in Fig G) and we computed the similarity between pRF profiles (Panel C in Fig G) and the cortical distance between every voxel of each ROI (Panel B in Fig G). We can see that for HC the similarity between pRF profiles is anticorrelated with the cortical distance and it is consistent across scanning sessions (t=0 $r^2$=-0.31; t=7d $r^2$=-0.29; t=17d $r^2$=-0.28; t=27d $r^2$=-0.28; Panel C in Fig G), showing that neighboring voxels have more similar pRF profiles than distant voxels. On the other hand, at t=0 for VD animals there is little to no correlation between the pRF profile and cortical distance (t=0 $r^2$=-0.14). With visual exposure, the VD pRF similarity coefficients become more similar to those of HC and progressively more anticorrelated with cortical distance (t=0 $r^2$=-0.14; t=7d $r^2$=-0.16; t=17d $r^2$=-0.2; t=72d $r^2$=-0.26; Panel C in Fig G). This approach further confirms the results of the phase gradient analysis and shows that light exposure drives a progressive reorganization of the visual pathway.

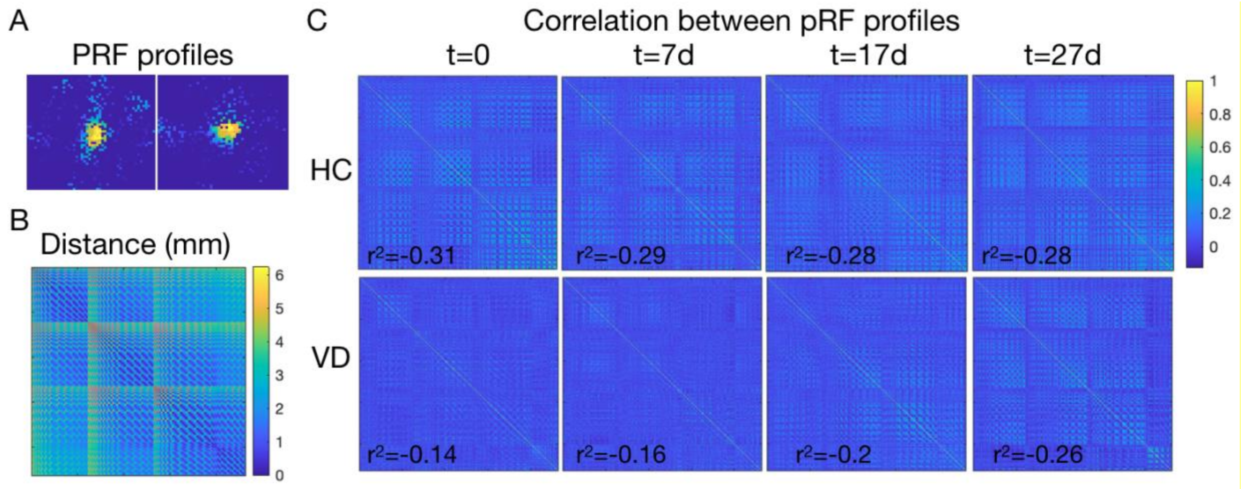


**Fig G. Quantification of the topographical organization of the visual pathway.** A: pRF profiles of 2 adjacent pRFs. B: Distance matrix of all the SC voxels. C: PRF similarity analysis of all SC voxels averaged for all the animals. The data underlying this figure can be found here: doi:10.18112/openneuro.ds004509.v1.0.0.

#### 7. Goodness of Gaussian fits to the SF tuning curves

| | Gaussian Fit Tuning Curve $r^2$ | | | | | | | |
|---|---|---|---|---|---|---|---|---|
| | HC | | | | VD | | | |
| | t=0 | t=7d | t=17d | t=27d | t=0 | t=7d | t=17d | t=27d |
| VC | 0.61 | 0.35 | 0.73 | 0.85 | 0.81 | 0.45 | 0.00 | 0.66 |
| LGN | 0.77 | 0.87 | 0.74 | 0.65 | 0.61 | 0.70 | 0.72 | 0.45 |
| SC | 0.68 | 0.27 | 0.85 | 0.88 | 0.70 | 0.89 | 0.83 | 0.59 |

**Table D. Pearson's coefficient between the maximum BOLD response to each SF and the Gaussian fit.** The data underlying this table can be found here: doi:10.18112/openneuro.ds004509.v1.0.0.

#### *8.* PRF variance explained in not visually responsive *brain areas*

We have calculated the pRF in the Auditory Cortex (AC), Motor Cortex (MC) and Basal Ganglia (BG) for the HC and VD animals at t=0. As expected the Variance Explained in the visual areas is higher than the VE in non-visual areas (~0.03). Note that in our analysis we have only considered the pRF whose variance explained is above 0.05. This excludes very noisy voxels and is well above the variance explained of the pRF is non-visual areas. The number of voxels excluded from the ROIs of the visual areas can be found in Table E. Moreover, the pRF profiles obtained for AC, MC and BG provide a good visualization of how noisy without a defined pRF are the measurements in areas that are not visually responsive (Fig H).

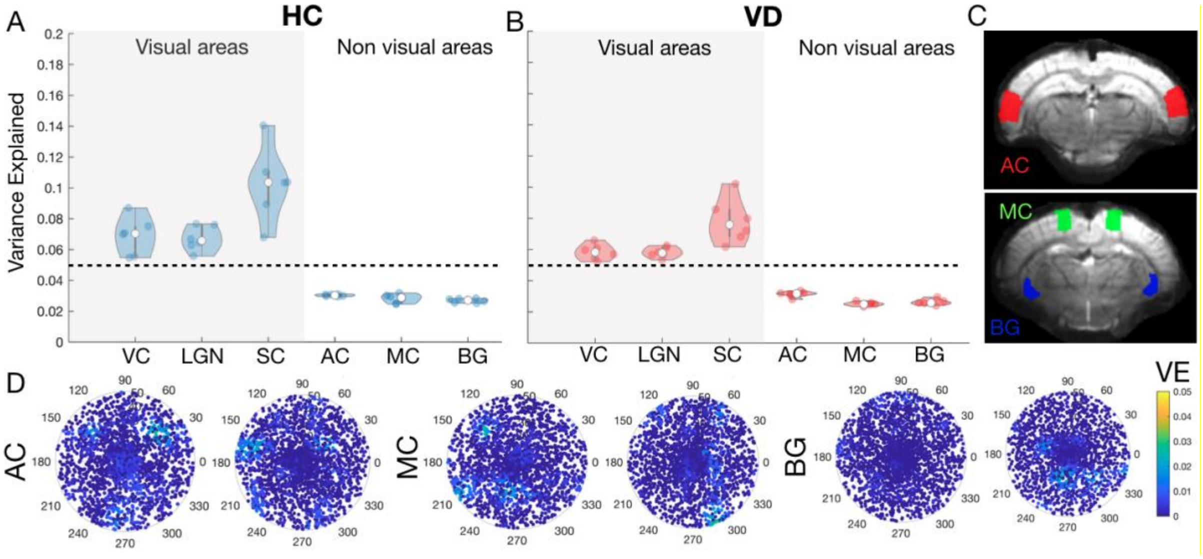


**Fig H. PRF estimates in the AC, MC and BG.** A and B: Violin plots of the variance explained calculated for visual areas (VC, LGN and SC) and for areas not visually responsive (AC, MC and BG) for HC and VD, respectively. C: Anatomical image with the ROIs AC, MC and BG overlapped. D: PRF profiles obtained for HC and VD animals (left and right, respectively) for AC, MC and BG. The data underlying this figure can be found here: doi:10.18112/openneuro.ds004509.v1.0.0.

| | HC | | | VD | | |
|---|---|---|---|---|---|---|
| | VC | LGN | SC | VC | LGN | SC |
| t=0 | 0.37±0.11 | 0.44±0.09 | 0.22±0.07 | 0.47±0.1 | 0.52±0.08 | 0.30±0.09 |
| t=7d | 0.50±0.12 | 0.54±0.12 | 0.30±0.09 | 0.47±0.07 | 0.52±0.06 | 0.3±0.06 |
| t=17d | 0.47±0.08 | 0.50±0.05 | 0.19±0.03 | 0.46±0.06 | 0.57±0.05 | 0.3±0.05 |
| t=27d | 0.42±0.12 | 0.45±0.09 | 0.23±0.12 | 0.47±0.06 | 0.56±0.07 | 0.22±0.04 |

**Table E. Fraction of voxels excluded by the variance explained threshold.** The data underlying this table can be found here: doi:10.18112/openneuro.ds004509.v1.0.0.

#### 9. The differences between HC and VD cannot be explained based on different vascular properties

In order to verify that the differences in BOLD amplitude and dynamics measured between HC and VD animals are not driven by different vascular properties between the two group of animals and between visual structures, the animals performed a hypercapnia challenge. A total of N = 5 HC (26 runs averaged) and N = 5 VD (27 runs averaged) were exposed to 1.5 min normocapnia, 1.5 min hypercapnia and 1.5 min normocapnia (Fig I). The rise times and signal amplitude have nearly identical onsets between VD and HC for the different ROIs (Panels B-G of Fig I). The mean PSC signal between VD and HC are highly correlated (VC: $r^2=0.95$ $p<0.001$; LGN: $r^2=0.86$ $p<0.001$; SC: $r^2=0.91$ $p<0.001$). In addition, a Granger causality test showed that HC hypercapnic response is very useful to predict the VD one (VC: F=10 p-value=0.0017; LGN: F=8.286 p-value=0.0045; SC: F=23 p-value=$3.27 \times 10^{-6}$). This excludes vasculature as the major contributing factor for the different measured timing parameters. The rise times and signal amplitude are nearly identical for all areas, suggesting that the vascular response dynamics becomes dissociated only at later stages.

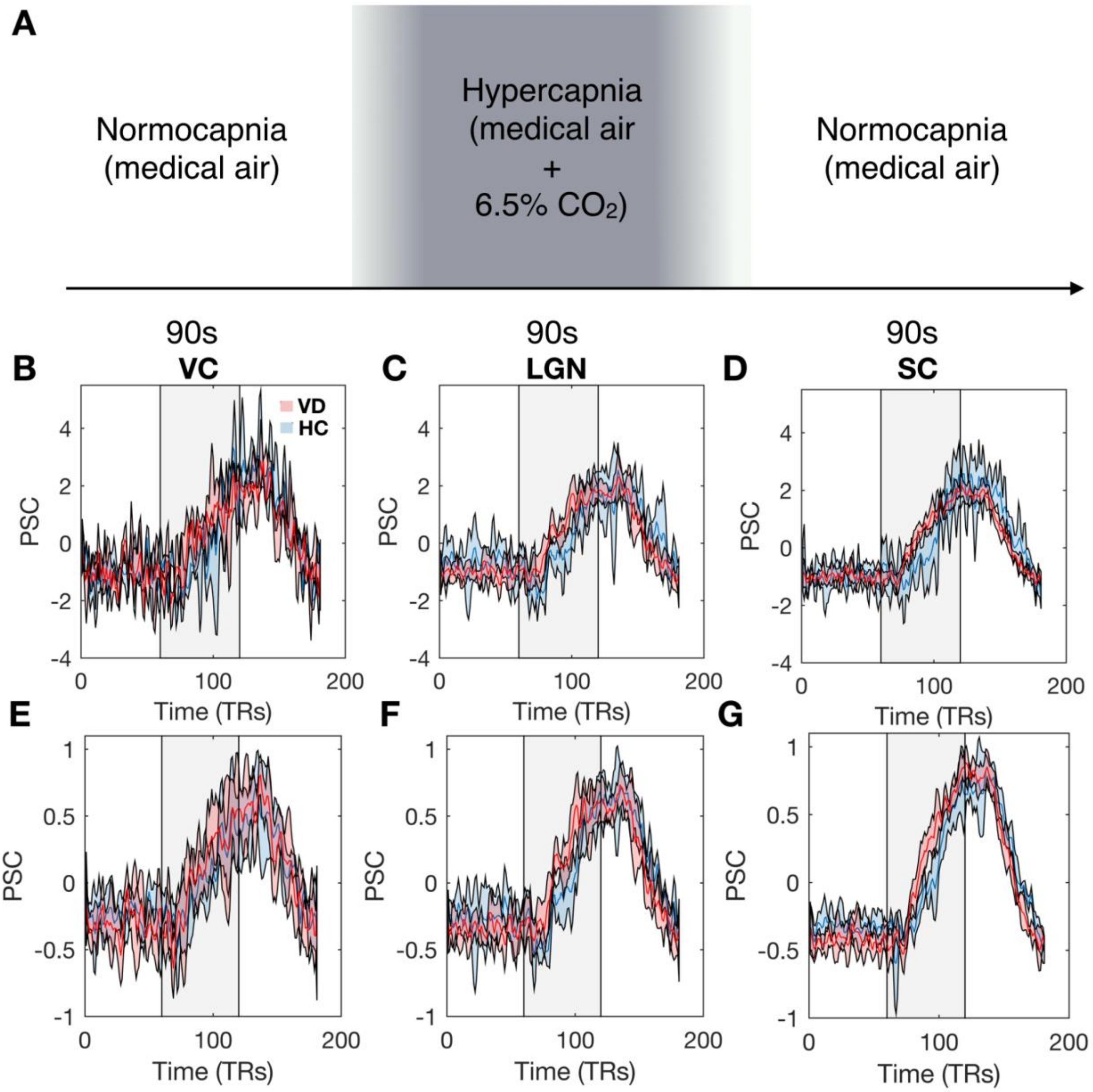


**Fig I. Hypercapnia experiment testing the dynamics of vascular responses.** A: Hypercapnia paradigm consisted of a manual switch, after 1.5 minutes of medical air, to a hypercapnic state with 6.5% $CO_2$ for 1.5 minutes. This was followed by a manual switch again to medical air for 1.5 minutes. Each run consisted in only one repetition of this block. B, C and D: PSC response profile (mean ± std) obtained for VD (red) and HC (blue) for different ROIs: VC, LGN and SC, respectively. The shaded grey area indicates the hypercapnic period. E, F and G: Normalized PSC response profile (mean ± std) obtained for VD (red) and HC (blue) for different ROIs: VC, LGN and SC, respectively. The data underlying this figure can be found here: doi:10.18112/openneuro.ds004509.v1.0.0.

### References:


1. Dodwell PC. Visual Orientation Preferences in the Rat. Quarterly Journal of Experimental Psychology.

1961. pp. 40–47. doi:10.1080/17470216108416467

19. Ito S, Feldheim DA, Litke AM. Segregation of Visual Response Properties in the Mouse Superior

Colliculus and Their Modulation during Locomotion. J Neurosci. 2017;37: 8428–8443.

20. De Franceschi G, Solomon SG. Visual response properties of neurons in the superficial layers of the superior colliculus of awake mouse. J Physiol. 2018;596: 6307–6332.

21. Prévost F, Lepore F, Guillemot J-P. Spatio-temporal receptive field properties of cells in the rat superior colliculus. Brain Res. 2007;1142: 80–91.
